## Supplementary for "Combining single-molecule super-resolved localization microscopy with fluorescence polarization imaging to study cellular processes"

### Supplementary figures and captions

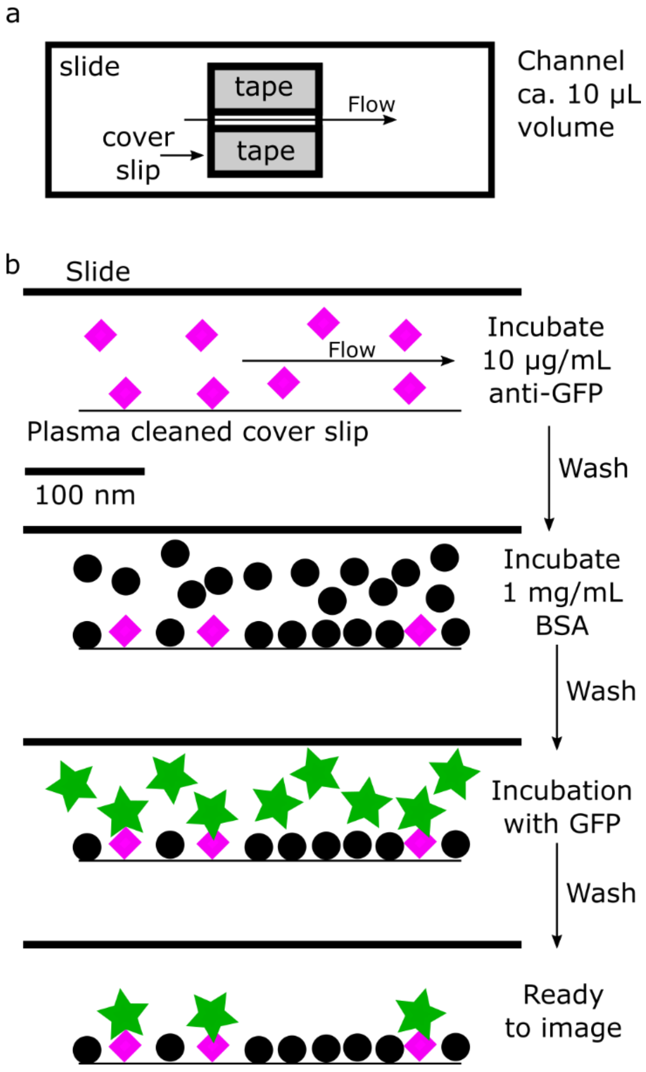


Supplementary Figure S1. a) A flow cell is created with a slide, plasma cleaned cover slip, and two lengths of double-sided tape. b) Schematic of the surface immobilized mGFP assay. First, anti-GFP is introduced to the flow cell and allowed to incubate. The anti-GFP (pink diamonds) has a high affinity for the plasma cleaned surface and are readily and strongly immobilized. After washing, 1 mg/mL BSA (black circles) is incubated to passivate the remaining exposed surface. After 5 minutes this is washed out and the mGFP itself (green stars) is introduced to bind with the anti-GFP antibody and allowed to incubate 5 minutes using a previously reported protocol [43]. After a final wash, the sample is ready to image (bottom panel). Before loading on to the sample, the channel can be hermetically sealed with nail polish or wax.

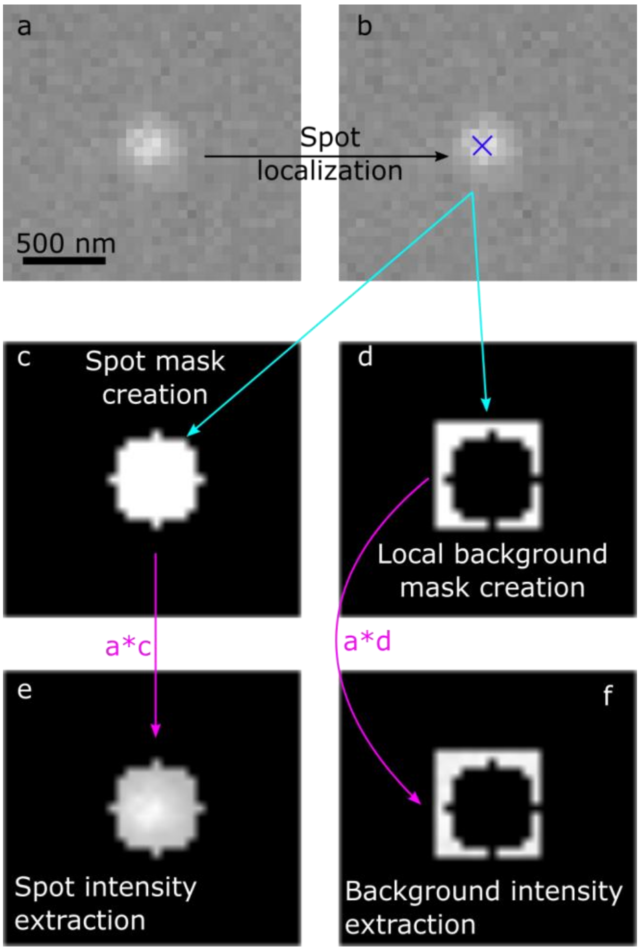


Supplementary Figure S2. a) Raw data from one of the polarization channels is passed to ADEMSCode (panel b) for spot fitting and subpixel localization. This localization is used to create two masks – one for the spot itself (c) and one for the local background (d). These are multiplied by the original image to give the intensities for each (e and f). The total fluorophore intensity is then the sum of each nonzero pixel in (e) after subtraction of the mean local background fluorescence in (f).
